## Supplemental information for "KAGE: Fast alignment-free graph-based genotyping of SNPs and short indels"

#### 1 Supplementary tables

| | Indels recall | Indels precision | Indels F1 | SNPs recall | SNPs precision | SNPs F1 | Runtime | Memory |
|---|---|---|---|---|---|---|---|---|
| KAGE | 0.587 | 0.912 | 0.714 | 0.936 | 0.982 | 0.959 | 12 min | 18 GB |
| KAGE + GLIMPSE | 0.594 | 0.911 | 0.719 | 0.948 | 0.989 | 0.968 | 1.4 hours | 18 GB |
| PanGenie | 0.566 | 0.938 | 0.706 | 0.912 | 0.987 | 0.948 | 4.4 hours | 113 GB |
| Bayestyper | 0.538 | 0.997 | 0.698 | 0.89 | 0.999 | 0.942 | 8.6 hours | 34 GB |
| Malva | 0.545 | 0.878 | 0.673 | 0.86 | 0.939 | 0.898 | 20.4 hours | 50 GB |
| Graphtyper | 0.592 | 0.938 | 0.726 | 0.941 | 0.994 | 0.967 | 11.6 hours | 33 GB |
| GATK | 0.97 | 0.98 | 0.975 | 0.982 | 0.983 | 0.983 | 19.0 hours | 67 GB |

**Supplementary Table 1. Results from genotyping HG002 at 30x coverage.**

| | Indels recall | Indels precision | Indels F1 | SNPs recall | SNPs precision | SNPs F1 | Runtime | Memory |
|---|---|---|---|---|---|---|---|---|
| KAGE | 0.628 | 0.922 | 0.747 | 0.914 | 0.981 | 0.946 | 12 min | 18 GB |
| KAGE + GLIMPSE | 0.638 | 0.91 | 0.75 | 0.932 | 0.991 | 0.96 | 1.3 hours | 18 GB |
| PanGenie | 0.605 | 0.94 | 0.736 | 0.889 | 0.979 | 0.932 | 4.0 hours | 113 GB |
| Bayestyper | 0.527 | 0.995 | 0.689 | 0.817 | 0.996 | 0.898 | 8.3 hours | 31 GB |
| Malva | 0.577 | 0.864 | 0.692 | 0.83 | 0.911 | 0.869 | 16.5 hours | 50 GB |
| Graphtyper | 0.61 | 0.957 | 0.745 | 0.889 | 0.993 | 0.938 | 5.9 hours | 20 GB |
| GATK | 0.905 | 0.96 | 0.932 | 0.947 | 0.977 | 0.962 | 9.6 hours | 58 GB |

**Supplementary Table 2. Results from genotyping HG006 at 15x coverage.**

| | Indels recall | Indels precision | Indels F1 | SNPs recall | SNPs precision | SNPs F1 | Runtime | Memory |
|---|---|---|---|---|---|---|---|---|
| KAGE | 0.633 | 0.923 | 0.751 | 0.921 | 0.978 | 0.949 | 22 min | 18 GB |
| KAGE + GLIMPSE | 0.639 | 0.923 | 0.755 | 0.932 | 0.985 | 0.958 | 1.5 hours | 18 GB |
| PanGenie | 0.609 | 0.949 | 0.742 | 0.894 | 0.986 | 0.937 | 4.5 hours | 113 GB |
| Bayestyper | 0.582 | 0.998 | 0.735 | 0.872 | 0.999 | 0.932 | 9.0 hours | 36 GB |
| Malva | 0.587 | 0.89 | 0.707 | 0.842 | 0.932 | 0.885 | 19.9 hours | 50 GB |
| Graphtyper | 0.635 | 0.955 | 0.763 | 0.922 | 0.995 | 0.957 | 10.1 hours | 34 GB |
| GATK | 0.967 | 0.977 | 0.972 | 0.966 | 0.977 | 0.972 | 17.5 hours | 58 GB |

**Supplementary Table 3. Results from genotyping HG006 at 30x coverage.**

#### 2 Details of method

##### 2.1 Problem Overview and Notation

We work with DNA sequences, represented as strings $s \in \Sigma^*$ over the alphabet

$$\Sigma = \{A, C, G, T\}$$

and variants over such strings: tuples of position, replaced sequence and replacing sequence that denote where and how a sequence differs from another sequence. For a reference sequence, and a variant, an individual can have genotype 0/0, 0/1 or 1/1 corresponding to whether that variant is present in none, one or both of the individual's genomic sequences (chromosomes). We let the set of variants be denoted $\mathcal{V} = N \times \Sigma^* \times \Sigma^*$, and the set of genotypes be denoted $\mathcal{G} = \{0/0, 0/1, 1/1\}$. Our goal is to predict where a diploid individual's genomic DNA sequence differs from a reference sequence (reference genome), and whether it differs in one or both of the individual's two haplotypes. As input to our algorithm we use:

- A reference genome sequence $\mathfrak{R} \in \Sigma^*$
- A list of $N$ known variants from the reference genome
- Information for a set of $M$ individuals on which genotype each individual have on each of the variants
- A set of sequenced reads from the individual of interest $\mathfrak{S} \in \Sigma^*$. These reads are assumed to be randomly sampled substrings from the individuals' full genomic sequence $I \in \Sigma^*$

Our method $F$ is then a function from these inputs to a list of predicted genotypes.

$$F : \Sigma^* \times V^N \times \mathcal{G}^{N \times M} \times \Sigma^* \rightarrow \mathcal{G}^N$$

The KAGE method can be divided into the following steps:

- Finding, for each variant, a representative kmer that is part of the reference sequence, but not part of the alternative sequence, and, vice versa, a kmer that is part of the alternative sequence but not part of the reference sequence.
- Finding, for each variant, another variant $v_h$ that correlates with our variant, i.e. where individuals tend to have the same genotype on the two variants. For each possible combination of genotypes on the two variants, calculate the probability of a random individual having that genotype combination.
- Count the number of occurrences of each kmer in the reference sequence (with variants).
- Count the number of occurrences of each kmer in the sequenced reads.
- For each variant, calculate the probability of getting the observed number of occurrences of the representative kmers for each of the possible genotypes.
- Combine the occurrence probabilities for each variant and its helper variant with the prior probabilities of having each genotype combination in a Bayesian formula to calculate the probability distribution over the genotypes.

From a computational perspective, it is important to note that step 1 through 3 is independent of the sequenced reads, and can thus be precalculated for a given reference genome and variant information (i.e GRCh38 and data from the 1000 Genomes Project).

##### 2.1.1 Kmer

Our model is mainly based on counting occurrences of kmers of length 31 (31-mers) in the different sequences. 31-mer are substrings of length 31, i.e $\in \Sigma^{31}$. We let $Occ(s, \mathfrak{X})$ denote the number of times kmer $s$ occurs in string $\mathfrak{X}$.

#### 2.2 Model formulation

Looking at a single variant of interest, we let $G_i$ denote the genotype on that variant, $G_h$ denote the genotype on the helper variant. We let $K_{ir}, K_{ia}$ denote the number of occurrences (in the sequenced reads) of the representative kmers for the reference and alternative sequence of the variant of interest, and similarly $K_{hr}$ and $K_{ha}$ for the helper variant. We want to calculate

$$\begin{aligned} P(G_i|K_{ir}, K_{ia}, K_{hr}, K_{ha}) &= \sum_{G_h \in \mathcal{G}} \frac{P(G_h, G_i, K_{hr}, K_{ha}, K_{ir}, K_{ia})}{P(K_{hr}, K_{ha}, K_{ir}, K_{ia})} \\ &= \sum_{G_h \in \mathcal{G}} \frac{P(G_h, G_i)P(K_{hr}, K_{ha}|G_h)P(K_{ir}, K_{ia}|G_i)}{P(K_{hr}, K_{ha}, K_{ir}, K_{ia})} \\ &\propto P(K_{ir}, K_{ia}|G_i) \sum_{G_h \in \mathcal{G}} P(G_h, G_i)P(K_{hr}, K_{ha}|G_h) \end{aligned}$$

The count probabilities $P(K_{ir}, K_{ia}|G_i)$ and $P(K_{hr}, K_{ha}|G_h)$ are calculated based on a Poisson model described in the next section, while the prior population probabilities $P(G_h, G_i)$ are based on the number of individuals having each combination of genotypes (described in the section Population Model).

#### 2.3 Count Model

We assume that the number of times a kmer $s$ occurs in the read set $\mathcal{S}$ is Poisson distributed with a rate proportional to the number of times $s$ occurs in the full genomic sequence of the sampled individual ($\mathfrak{I} \in \Sigma^*$). I.e. $Occ(s, \mathfrak{S}) \sim \text{Pois}(\lambda_0 Occ(s, \mathfrak{I}))$. Since we don't know the exact sequence of the sampled individual (this is in part what we are trying to find out), we can't calculate this probability directly. Based on the reference genome and the variant information, we can however deduce an approximate probability distribution over $Occ(s, \mathfrak{I})$. Based on this we can calculate:

$$P(Occ(s, \mathfrak{S}) = k) = \sum_{d=0}^{\infty} P(Occ(s, \mathfrak{I}) = d)\text{Pois}(k; \lambda_0 d)$$

.

##### 2.3.1 Estimating the occurrence distribution

How many times a kmer occurs in an individuals genome sequence can depend on which variants the individual has. In order to account for this, we use genotype information from a reference population, like the 1000 Genomes Project, to estimate the probabilities $P(occ(s_r, \mathfrak{I}) = d|G = g))$ by simply counting the number of individuals with a given genotype that has $d$ occurrences of the kmer. Using these occurrence probabilities we can calculate the the probabilities for observed counts given a genotype as:

$$P(K_a|G) = \sum_{d=0}^{\infty} P(Occ(s_a, \mathfrak{I}) = d|G)\text{Pois}(\lambda_0(d+\epsilon))$$
$$P(K_r|G) = \sum_{d=0}^{\infty} P(Occ(s_r, \mathfrak{I}) = d|G)\text{Pois}(\lambda_0(d+\epsilon))$$

### 3 Population Model

As input to the method we get information for each variant on which individuals have which genotype on that variant. For a pair of variant of interest and helper variant, we can thus calculate the number of individuals that have a given combination of genotypes $(G_i, G_h)$, denoted $\pi(G_i, G_h)$. We use these counts to infer the conditional probabilities $P(G_i|G_h)$ by assuming a Dirichlet prior over the genotypes according to the average number of individuals having the different combinations over all interest-helper variant pairs. We set this prior with a concentration $\alpha_0 = 5$. Thus we get that

$$P(G_i|G_h) = \frac{\alpha_0 G_i|G_h + \pi(G_i, G_h)}{\sum_{g_i \in \mathcal{G}} \alpha_{g_i|G_h} + \pi(g_i, G_h)}$$

where

$$\alpha_{G_i|G_h} = \alpha_0 \frac{\sum_v 1 + \pi_v(G_i, G_h)}{\sum_{g_i \in \mathcal{G}} \sum_v 1 + \pi_v(g_i, G_h)}$$

Similarly, we find $\mathrm{P(G_h)}$ by using a Dirichlet prior according to the average across all variants. I.e.

$$P(G_h) = \frac{\beta_{G_h} + \sum_{G_i \in \mathcal{G}} \pi(G_i, G_h)}{\sum_{g_h \in \mathcal{G}} \beta_{g_h} + \sum_{G_i \in \mathcal{G}} \pi(G_i, g_h)}$$

where

$$\beta_{G_h} = \beta_0 \frac{\sum_v 1 + \sum_{G_i \in \mathcal{G}} \pi_v(G_i, G_h)}{\sum_v \sum_{g_h \in \mathcal{G}} 1 + \sum_{G_i \in \mathcal{G}} \pi_v(G_i, G_h)}$$

In less formal terms, this corresponds to adding weighted pseudo counts to the observed population counts. To get the probability distribution over genotype combinations, we simply multiply these:

$$P(G_i, G_h) = P(G_h)P(G_i|G_h)$$

These values are precalculated for each variant pair.

#### 3.1 Finding the helper variant.

For each variant of interest, we want to find one other variant that correlates well with our variant of interest. This means that for a good helper variant, individuals tend to have the same genotype on the helper variant as on the variant of interest. Using such variants we can improve our prediction of the interest-genotype in cases where the count model for the variant of interest gives ambiguous results. In order to find good helper variants, we evaluate the metric $\sum_{G \in \mathcal{G}} \log \pi_v(G, G)$ for the neighbouring 200 variants to the variant of interest, and choose as helper the variant $v$ with the highest score.
